# Heterogeneity and genomic loci of ubiquitous Cre reporter transgenes in zebrafish

**DOI:** 10.1101/2021.12.22.473906

**Authors:** Robert L. Lalonde, Cassie L. Kemmler, Fréderike W. Riemslagh, Andrew J. Aman, Jelena Kresoja-Rakic, Hannah R. Moran, Susan Nieuwenhuize, David M. Parichy, Alexa Burger, Christian Mosimann

## Abstract

The most-common strategy for zebrafish Cre/*lox*-mediated lineage labeling experiments combines ubiquitously expressed, *lox*-based *Switch* reporter transgenes with tissue-specific Cre or 4-OH-Tamoxifen-inducible CreERT2 driver lines. Although numerous Cre driver lines have been produced, only a few broadly expressed Switch reporters exist in zebrafish and their generation by random transgene integration has been challenging due to position-effect sensitivity of the *lox*-flanked recombination cassettes. Here, we compare commonly used *Switch* reporter lines for their recombination efficiency and reporter expression pattern during zebrafish development. Using different experimental setups, we show that *ubi:Switch* and *hsp70l:Switch* outperform current generations of two additional *Switch* reporters due to favorable transgene integration sites. Our comparisons also document preferential Cre-dependent recombination of *ubi:Switch* and *hsp70l:Switch* in distinct zebrafish tissues at early developmental stages. To investigate what genomic features may influence Cre accessibility and *lox* recombination efficiency in highly functional *Switch* lines, we mapped these transgenes and charted chromatin dynamics at their integration sites. Our data documents the heterogeneity among *lox*-based *Switch* transgenes towards informing suitable transgene selection for lineage labeling experiments. Our work further proposes that *ubi:Switch* and *hsp70l:Switch* define genomic integration sites suitable for universal transgene or switch reporter knock-in in zebrafish.

## Introduction

Site-specific recombinase (SSR)-based techniques provide powerful versatility to transgenic models. Techniques based on Cre/*lox*, Flp/*FRT*, or phiC31 recombinase systems allow permanent excision or rearrangement of transgene cassettes to modify their activity and function (Branda and Dymecki, 2004; Carney and Mosimann, 2018; Guillou, 2006; McLellan et al., 2017; Rossant and Nagy, 1995). As widely used recombinase system in vertebrate models, the Cre/*lox* system combines i) a Cre recombinase-providing transgene driven by a tissue-specific *cis-*regulatory element and ii) a recombination-competent reporter transgene. The Cre recombinase, originally derived from the P1 bacteriophage, directionally recombines DNA at specific 13 bp palindromic repeats called *loxP* sites in a variety of species including mice and zebrafish (Branda and Dymecki, 2004; Langenau et al., 2005; Rossant and Nagy, 1995; Sauer, 1987). A major application of the Cre/*lox* system in zebrafish involves the recombination of *lox*-flanked fluorophore cassettes that switch upon Cre activity to lineage-label cell populations of interest. This strategy has been successfully employed to reveal developmental lineage origins, to follow post-embryonic stem cells, and to track the lineage composition of regenerating organs (Carney and Mosimann, 2018; Choi et al., 2014; Dirian et al., 2014; F et al., 2011; Felker et al., 2018; Gupta et al., 2013; Kaufman et al., 2016; Le et al., 2007; Li et al., 2019; Mosimann et al., 2011). These experiments hinge upon combining a tissue- or cell type-specific Cre or 4-OHT-Tamoxifen (4-OHT)-inducible CreERT2 driver with temporal control and a broadly active or ubiquitous *lox*-based reporter transgenic (Feil et al., 1997; Hans et al., 2009; Hans et al., 2011; Mosimann et al., 2011). Particularly simple to control in developing zebrafish by 4-OHT addition to the embryo medium, CreERT2 transgenics have rapidly increased in number across the field and include Tol2-based transgenes driving CreERT2 with tissue-specific regulatory elements, gene trap collections, and first CRISPR-based knock-in lines (Carney and Mosimann, 2018; Hans et al., 2009; Jungke et al., 2013; Jungke et al., 2015; Kesavan et al., 2018). In contrast, the generation of suitable and reproducibly well-performing *lox*-based reporter transgenes has proven challenging.

The heat shock protein 70-like (*hsp70l*), *beta-actin2* (*actb2)*, and *ubiquitin* (*ubi* or *ubb*) promoter elements have been routinely used to drive quasi-ubiquitous transgene gene expression in zebrafish and have been successfully applied in several *lox*-based *Switch* reporters (Blechinger et al., 2002; Burket et al., 2008; Carney and Mosimann, 2018; Felker et al., 2018; Higashijima et al., 1997; Kobayashi et al., 2014; Langenau et al., 2005; Mosimann et al., 2011). *hsp70l*-based transgenes are a mainstay of the zebrafish’s transgenic tool kit, as heat shock induction around 37°C causes rapid ubiquitous reporter expression that is however not sustained long beyond the heat shock pulse (Blechinger et al., 2002). The *actb2* promoter provides strong, rapid transgene expression during development and in individual tissues, yet its activity drastically diminishes or silences in select cell types such as erythrocytes (Burket et al., 2008; Chen et al., 2017; Higashijima et al., 1997; Traver et al., 2003). The *ubi* promoter drives widespread and persistent transgene expression in various independent transgenics including the Cre-sensitive *loxP* reporter *ubi:loxP-GFP-loxP_mCherry* (*ubi:Switch*) that has found widespread use (Mosimann et al., 2011). Nonetheless, *ubi*-based transgenics including *ubi:Switch* show slow reporter accumulation at early developmental stages (Chen et al., 2017; Felker et al., 2016; Mosimann et al., 2011); this property causes considerable latency between Cre-triggered *lox* cassette recombination and reporter detection, rendering short lineage labeling timeframes (under 24 hours) challenging to achieve.

The advantages and drawbacks of each regulatory element require careful characterization of individual transgene insertions over multiple generations, data that is rarely available to guide experimental design. Generating reproducibly functional, single-insertion *lox-*based *Switch* transgenics by randomly integrating Tol2 or ISce-I transgenesis is also notoriously screening-intensive due to position-effect sensitivity of *lox* cassette recombination (Carney and Mosimann, 2018; Kawakami et al., 2004; Kikuta and Kawakami, 2009). Consequently, together with the challenges of sharing transgenic zebrafish lines internationally, the majority of labs only have access to one or few Switch reporter lines. Means to efficiently generate Switch transgenics are therefore highly desirable, requiring the identification of suitable genomic loci for transgene knock-ins.

Here, we compared previously published and validated *Switch* reporter lines in combination with ubiquitous and tissue-specific CreERT2 drivers. Our results document the heterogeneity in recombination efficiency and preferential tissue expression of individual reporters. Together with genomic integration mapping of tested transgenes, our results define *ubi:Switch and hsp70l:Switch* as integrations in favorable loci for *lox* cassette recombination and transgene expression.

## Results

### Ubiquitous *Switch* reporters feature variable recombination efficiency and tissue activity

To gain insight into recombination efficiency and reporter expression across tissues by ubiquitous *Switch* reporters in development, we compared transgenic zebrafish reporters that have been previously used to follow lineage trajectories. We focused on *Switch* lines in our collection that have been documented to drive ubiquitous transgene expression: *ubi:Switch*^*cz1701*^, *hsp70l:Switch*^*zh701*^, *Tg(bactin2:loxP-Stop-LoxP-DsRed-express)*^*sd5*^ (shortened to *actb2:Stop-DsRed*), and *Tg(bactin2:loxP-BFP-loxP-DsRed)*^*sd27*^ (shortened to *actb2:BFP-DsRed*) (Bertrand et al., 2010; Felker et al., 2018; Kobayashi et al., 2014; Mosimann et al., 2011). The *ubi:creERT2* driver has been widely used for ubiquitous CreERT2 expression in early development and provides a first assay for Cre responsiveness of individual *Switch* lines (Mosimann et al., 2011). To establish switching efficiency under *ubi:creERT2* and homogeneity versus heterogeneity of reporter expression, we treated *ubi:creERT2;promoter:Switch* embryos with 4-OHT at shield stage and imaged whole larvae at 3 dpf (**Fig. 1**). In all crosses, the *ubi:creERT2*-carrying parent was male to avoid maternal CreERT2 contribution.

**Figure 1.**
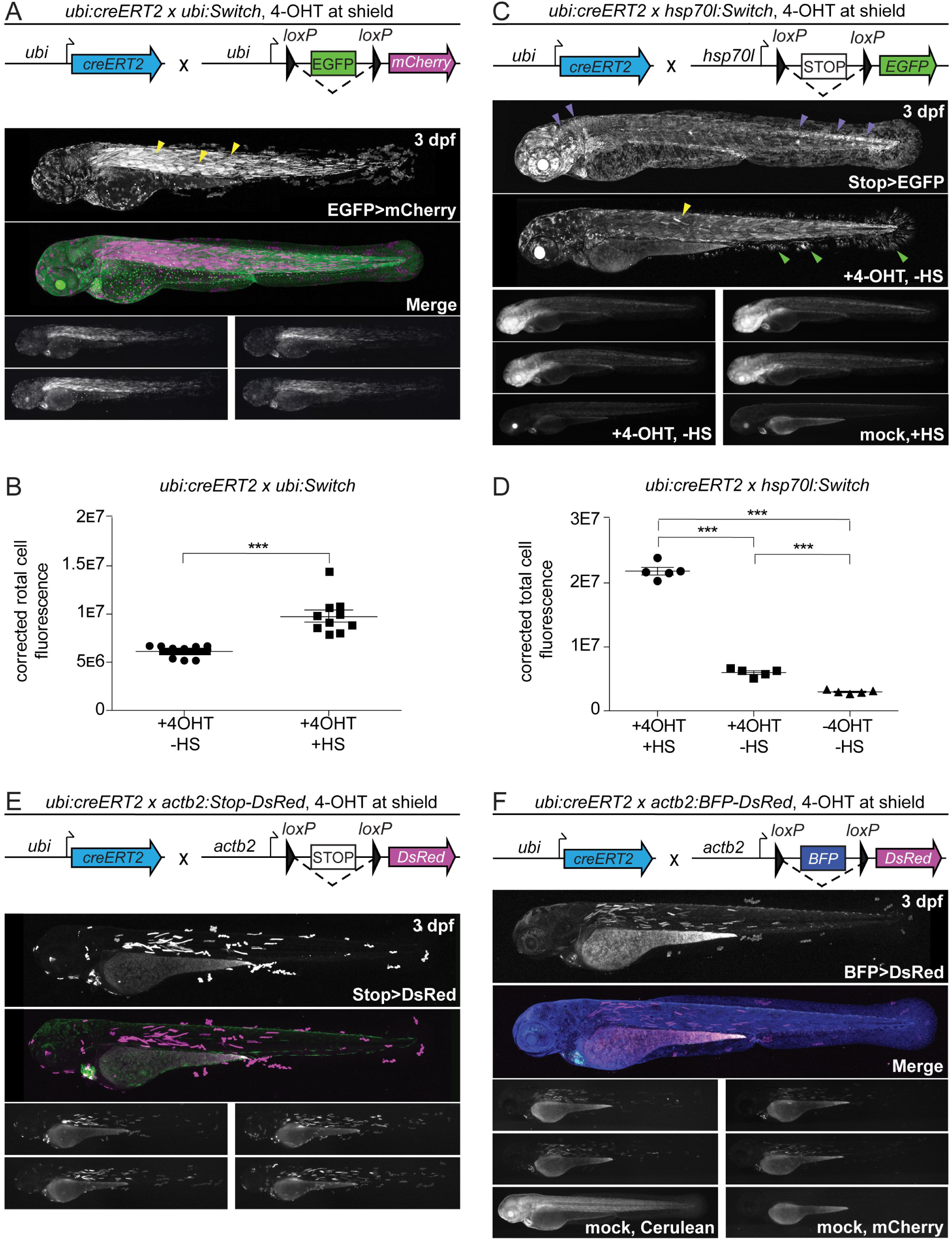
Ubiquitous *Switch* reporter lines show variable recombination efficiency. (**A**,**B**) *ubi:Switch*, (**C**,**D**) *hsp70l:Switch*, (**E**) *actb2:Stop-DsRed*, and (**F**) *actb2:BFP-DsRed* crossed to *ubi:creERT2*, induced with 10 µM 4-OHT at shield stage, and imaged laterally at 3dpf. Schematics of fluorophore cassettes for each *Switch* transgene are shown at the top of each panel (**A**,**B**,**E**,**F**). One representative confocal image, and four representative stereo microscope images are presented per reporter. *ubi:Switch* (**A**) shows preferential recombination in somitic muscle (yellow arrowheads), *hsp70l:Switch* (**C**) shows preferential switching in CNS (brain, neural tube) (purple arrowheads). Non heat-shocked (+4-OHT) controls (**C**, second panel) show faint EGFP expression in somitic muscle (yellow arrowheads) and fin fibroblasts (green arrowheads). (**B**) Corrected total cell fluorescence (CTCF) measurements of 3dpf *ubi:creERT2* crossed to *ubi:Switch*, induced with 10 µM 4-OHT at shield, and heat-shocked 3 hours prior to lateral view imaging; fluorescence intensity was compared to non-heat-shocked sibling (n=10, 2 clutches). Note increased CTCF following heat-shock at 3 dpf (Mann-Whitney, P>0.0001). (**B**) CTCF *ubi:creERT2* crossed to *hsp70l:Switch*, induced with 10 µM 4-OHT at shield. Non-heat shocked controls (center), and non-heat shocked, non-treated controls (right) are included (1-way ANOVA, P<0.0001). (**E**,**F**) *ubi:Switch* and *hsp70l:Switch* show more spatially complete recombination compared to both *actb2:Stop-dsRED* (**E**) *and actb2:BFP-DsRed* (**F**) lines that only display sparse recombination.

The *ubi:Switch* transgenic permanently recombines from EGFP to mCherry following Cre activity (**Fig. 1A**), yet mCherry levels only reach detectable levels after several hours or a day post-Cre activity (Mosimann et al., 2011). When combined with *ubi:creERT2*, we observed strongest mCherry activity in the somitic muscle along the entire trunk and tail (**Fig. 1A**). Skin and fin epithelium showed more sparse switching, predominantly in the median fin and head (**Fig. 1A**). We observed strong mCherry signal in the heart, while other tissues including the eye lens and neural tube consistently displayed comparatively lower mCherry fluorescence (**Fig. 1A**). Promoters of *ubiquitin* genes in several species have been postulated to harbor heat shock-responsive elements, possibly to support protein degradation by increased *Ubiquitin* polypeptide production upon heat or other stress (Bond and Schlesinger, 1985; Christensen et al., 1992; Fornace et al., 1989; Fujimuro et al., 2005; Lee et al., 1988; Nenoi et al., 1996). To test if *ubi:Switch* as driven by the zebrafish *ubb* gene promoter reaches higher expression levels upon heat shock, we compared the levels of mCherry fluorescence at 3 dpf with and without a 1 hour-long, 37°C heat shock prior to imaging: compared to non-heat-shocked siblings, heat-shocked *ubi:creERT2;ubi:Switch* larvae showed increased fluorescent intensity 3 hours post-heat shock (n = 10 for quantification, 2 clutches) (**Fig. 1B**). These observations document that *ubi:Switch* expression can be further augmented by a brief heat shock prior to imaging.

Following Cre activity, the *hsp70l:Switch* transgenic line permanently switches from a non-fluorescent Stop cassette to EGFP (Felker et al., 2018). Heat shock for 1 hour at 37°C induces prominent EGFP expression within one hour post-heat shock treatment. To trigger reporter expression of *hsp70l:Switch*, we heat-shocked recombined embryos 2-3 hours prior to imaging (**Fig. 1**). When crossed to *ubi:creERT2* and treated with 4-OHT at shield stage, *hsp70l:Switch* showed seemingly uniform switching across the entire embryo, with the central nervous system (brain, neural tube) displaying higher EGFP signal (**Fig. 1B**). Notably, in non-heat-shocked controls that were 4-OHT-induced, we also detected faint EGFP expression (n = 5) in the skeletal muscle and fin fibroblasts (**Fig. 1C,D**); these observations indicate that tissue-specific accessibility of the *hsp70l* element at the genomic location of *hsp70l:Switch* leads to transcriptional activity even at 28°C. Embryos that were left untreated with 4-OHT did not show any detectable EGFP expression with or without heat shock, confirming that the used stop cassette (Felker et al., 2018; Hesselson et al., 2009) efficiently interrupts any EGFP transcription.

The *actb2:Stop-DsRed* transgene recombines from no fluorophore to DsRed and *actb2:BFP-DsRed* recombines from BFP to DsRed, enabling versatile combination with existing tissue-specific reporters based on EGFP (Bertrand et al., 2010; Kobayashi et al., 2014). After recombination with *ubi:creERT2* at shield stage, both *actb2 Switch* lines imaged at 3 dpf showed overall sporadic switching with the most prominent DsRed signal in the skeletal muscle, fin epidermis, and the heart (**Fig. 1E,F**). Under these conditions, switching was absent from a large majority of tissues including, but not limited to, endothelial cells, skin, brain and neural tube, and fin fibroblasts (**Fig. 1E,F**).

Taken together, our observations from triggering ubiquitous CreERT2 activity after onset of gastrulation reveal heterogeneous recombination efficiencies and reporter expression across four distinct, broadly expressed *lox* reporter lines. The results with *ubi:Switch* are in line with previously reported Cre-based lineage labeling experiments across various cell types and developmental stages. Nonetheless, *hsp70l:Switch* displayed distinctively more homogenous reporter expression than *ubi:Switch*; this effect is possibly in part due to the pulsed fluorescence activation following heat shock shortly before analysis in *hsp70l:Switch* experiments, while in *ubi:Switch* the mCherry reporter dynamically accumulates and degrades in various cell types following 4-OHT treatment. In contrast, the lower switching efficiency observed with the *actb2-*based *Switch* lines indicates low Cre accessibility of the insertions selected for these transgenics over the past decade (Bertrand et al., 2010; Kobayashi et al., 2014).

### *hsp70l:Switch* shows widespread and reproducible Cre sensitivity

The widespread Cre-induced recombination and reporter expression observed with *ubi:Switch* and *hsp70l:Switch* indicate favorable genomic integration sites for *loxP* cassette recombination. To further evaluate the switching efficiencies between *ubi:Switch* and *hsp70l:Switch*, we turned to two previously established, highly active, and tissue-specific CreERT2 driver lines: i) *Tg(drl:creERT2;cryaa:Venus)* (*drl:creERT2* for short) that expresses in lateral plate mesoderm (LPM)-primed mesendoderm during gastrulation before refining to cardiovascular lineages by mid-somitogenesis (Mosimann et al., 2015; Prummel et al., 2019) (**Fig. 2A-C**); and ii) *Tg(tbx1:creERT2;cryaa:Venus)* (*tbx1:creERT2* for short) that starts expressing in late gastrulation and predominantly labels cardiopharyngeal progenitors together with *tbx1*-expressing ectodermal and endodermal lineages in the head (Felker et al., 2018) (**Fig. 2D-F**). After crossing, we treated double-transgenic embryos with 10 µM 4-OHT at shield stage and imaged the head region at 3 dpf laterally and ventrally to assess lineage labeling.

**Figure 2.**
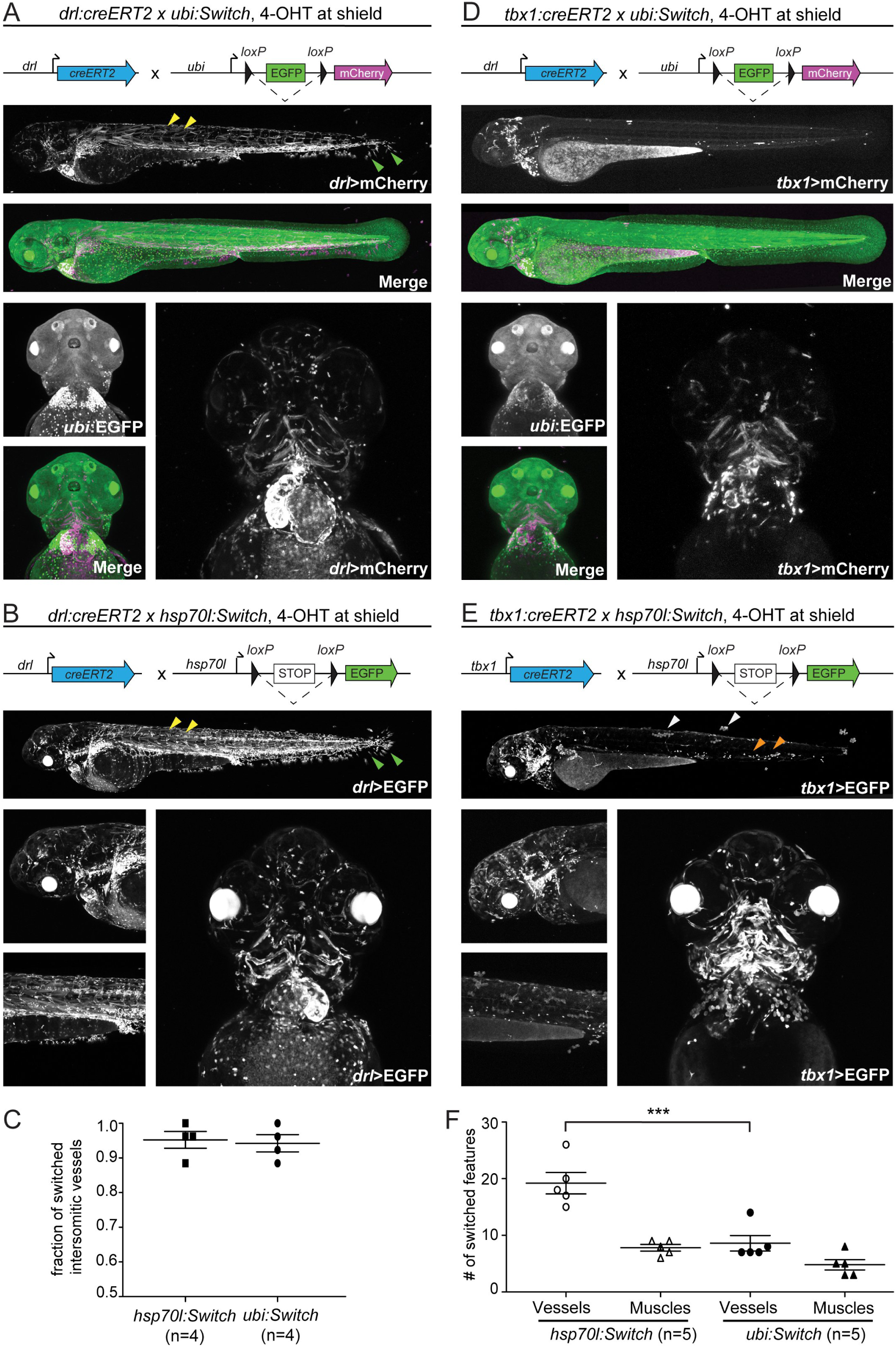
*hsp70l:Switch* high degrees of lineage labeling crossed to tissue-specific CreERT2 driver lines. (**A**,**B**) *drl:creERT2*, and (**C**,**D**) *tbx1:creERT2* crossed to *ubi:Switch* and *hsp70l:Switch*, respectively induced with 4-OHT at shield stage and imaged laterally and ventrally at 3dpf. *drl:creERT2* yields more complete recombination when crossed to *hsp70l:Switch* (**B**) compared to *ubi:Switch* (**A**). The higher degree of recombination is detectable in the somitic muscle (yellow arrowheads), and fin fibroblasts (green arrowhead) (**A**,**B**). No difference is observed in percentage of switched intersomitic vessels (ISV) when *drl:creERT2* is crossed to either *hsp70l:Switch* or *ubi:Switch* (Mann-Whitney, p>0.05) (**C**). *tbx1:creERT2* yields less switching mosaicism when crossed to *hsp70l:Switch* (**E**), compared to *ubi:Switch* (**D**). Higher recombination efficiency is readily observed in trunk skin (white arrowhead) and hematopoietic cells (orange arrowheads) (**D**,**E**). When crossed to *hsp70l:Switch, tbx1:creERT2* yields significantly more switched head vessels compared to *ubi:Switch* (1-way ANOVA p<0.0001) (**F**). Representative lateral and ventral images (**A**,**B**;**D**,**E**).

Combined with *drl:creERT2*, both *Switch* lines displayed switching in the same tissues and cell types, yet *hsp70l:Switch* consistently resulted in more complete recombination (lower mosaicism). This feature was most evident in the prevalence of switched fin fibroblasts, skeletal muscle cells, macrophages, and head vasculature (**Fig. 2A,B**). To compare recombination efficiency in a major cell type, we quantified switched versus unlabeled intersomitic vessels (ISV) (Felker et al., 2016): we observed no significant difference in percentage of switched ISV between *ubi:Switch* or *hsp70l:Switch* when recombined by *drl:creERT2* (n = 4 for quantification) (**Fig. 2C**). Of note, induction of *drl*:*creERT2* at shield stage results in complete labeling of LPM-derived structures, but also sparse labeling of paraxial mesoderm-derived structures including somitic skeletal muscle cells and median fin fold fibroblasts, reflecting the starting segregation of individual mesodermal territories at shield stage (Mosimann et al., 2015; Prummel et al., 2019) (**Fig. 2A,B**).

When crossed to *tbx1:creERT*, both *ubi:Switch* and *hsp70l:Switch* displayed broad lineage labeling in the ventricular cardiomyocytes, pharyngeal arches, cranial vasculature, head muscles and cartilage, and hatching gland, consistent with previous work (Felker et al., 2018) (**Fig. 2D,E**). Once more, *ubi:Switch* animals showed preferential switching in muscles, versus *hsp70l:Switch* that displayed more complete switching across multiple cell types, especially head vasculature. We quantified muscle and endothelial switching of each *Switch* line using muscle (MF20 antibody) and endothelial (Fli1 antibody) staining on recombined larvae compared to the endothelial reporter *fli1:EGFP* as reference (n=5) (**Fig. 2F, Fig. S1**). *tbx1:creERT2* had significantly higher recombination efficiency in the head vasculature combined with *hsp70l:Switch* than when combined with *ubi:Switch*, but with no significant difference in head muscle switching between the two *Switch* lines (**Fig. 2F**). These results further underline the widespread, yet differential sensitivity to Cre activity of both *ubi:Switch* and *hsp70l:Switch*.

### *ubi:creERT2* shows decreased recombination activity at 48 hpf

The *ubi:creERT2*^*cz1702*^ transgenic line has been widely applied to provide broad 4-OHT-inducible Cre activity, as desirable for testing new *Switch* transgenes (Carney and Mosimann, 2018; Mosimann et al., 2011). Our data presented above further confirmed the reproducible activity of *ubi:creERT2* at shield stage paired with responsive *Switch* lines. Nevertheless, anecdotal observations have suggested that *ubi:creERT2* becomes less responsive to 4-OHT at later stages (Felker et al., 2016; Mosimann et al., 2011). To what extent this effect depends on choice of *lox Switch* line remains uncertain.

To test the recombination capacity after gastrulation stages, we crossed *ubi:creERT2* to *ubi:Switch* and *hsp70l:Switch*, treated with 10 μM 4-OHT at 48 hpf, and performed lateral-whole embryo imaging at 3dpf (**Fig. 3**). In contrast to shield stage induction, 4-OHT induction at 48 hpf showed markedly reduced switching at 3 dpf with both *ubi:Switch* and *hsp70l:Switch* (**Fig. 3A,B**). When crossed to *hsp70l:Switch*, 10 μM 4-OHT induction at 48 hpf of *ubi:creERT2* resulted in switching predominantly in the skeletal muscle and the fin fibroblasts **(Fig. 3A)**. When crossed to *ubi:Switch* and induced with this regimen, *ubi:creERT2* induced switching predominantly in the blood and heart (**Fig. 3B**). Due to the cardiac *myl7:EGFP* reporter incorporated in the *ubi:creERT2* transgene, we could not evaluate switching efficiency in the heart using *hsp70l:Switch*. While 10 μM results in reproducible CreERT2 activity across transgenes with negligible impact on the treated embryos, previous work has indicated that 4-OHT reaches saturation for inducing CreERT2 activity at 25 μM (Felker et al., 2016; Hans et al., 2011; Mosimann et al., 2011). In line with this observation, increasing 4-OHT concentration from 10 μM to 30 μM at 48 hpf did not notably increase switching efficiency in *ubi:creERT2*;*ubi:Switch* embryos (**Fig. S1**).

**Figure 3.**
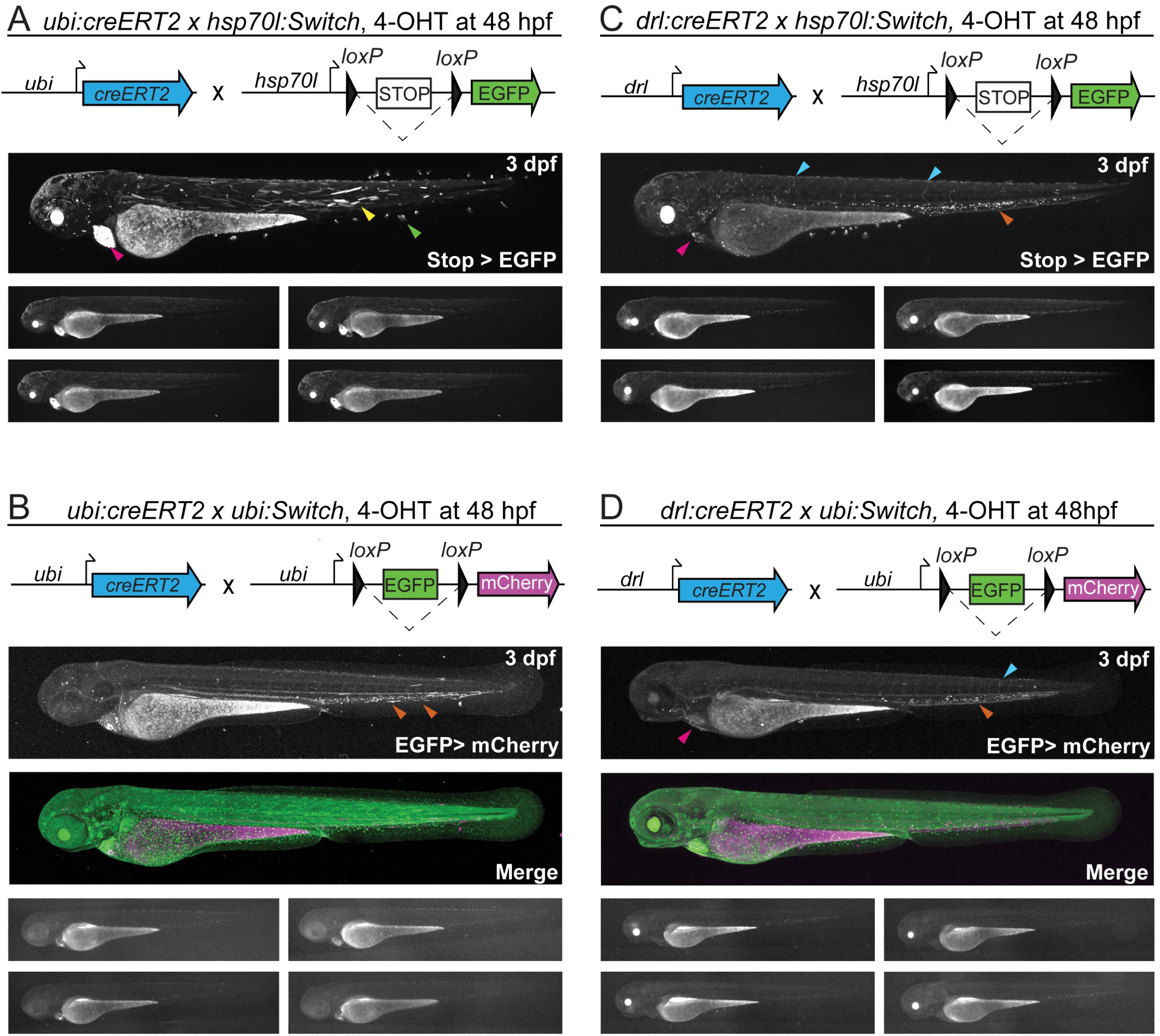
*ubi:creERT2* shows decreased 4-OHT and recombination response at larval stages. (**A**,**B**) *ubi:creERT2* and (**C, D**) *drl:creERT2* crossed to *ubi:Switch*, and *hsp70l:Switch*, induced with 4-OHT at 48hpf, and imaged laterally at 3dpf. Schematics of fluorophore cassettes for each Switch transgene are shown at the top of each panel; one representative confocal image, and four representative stereo microscope images are presented here (**A-D**). *ubi:creERT2* shows sparse switching with both reporters when induced with 4-OHT at 48 hpf (**A**,**B**). Predominant switching occurs in somitic muscle (yellow arrowheads), and fin fibroblasts (green arrowheads) when crossed to *hsp70l:Switch* (**A**), and hematopoietic cells (orange arrowhead) when crossed to *ubi:Switch* (**B**). *drl:creERT2* with 4-OHT induction at 48 hpf results in switching in the heart (pink arrowhead), vasculature (blue arrowhead), and hematopoietic cells (orange arrowhead) when crossed to both *hsp70l:Switch* (**C**), and *ubi:Switch* (**D**). Note for *ubi:Switch* stereo microscope images that are representative of routine laboratory imaging, extended exposure (30 seconds) was used to capture all traces of mCherry fluorescence (**B**,**D**).

To test whether the observed decrease in switching efficiency was specifically associated with the *ubi:creERT2* line, we crossed *drl:creERT2* to *hsp70l:Switch* and *ubi:Switch*, treated with 10 μM 4-OHT at 48 hpf, and performed lateral-whole embryo imaging at 3 dpf (**Fig 3C,D)**. *drl*-based transgenics is restricted predominantly to cardiovascular and hematopoietic lineages by 48 hpf (Mosimann et al., 2015; Prummel et al., 2019). Accordingly, when crossed to *hsp70l:Switch*, 48 hpf induction of *drl:creERT2* resulted in robust switching in hematopoietic cells with sporadic labeling in the heart, endothelial cells, as well as in pectoral fin fibroblasts (**Fig. 3C)**. When crossed to *ubi:Switch*, 48 hpf induction of *drl:creERT2* again resulted in robust switching in hematopoietic cells, with sporadic labeling in the heart, endothelial cells, and fin fold fibroblasts (**Fig. 3D)**. Despite maximum exposure lengths, visualizing mCherry fluorescence at 24 hours post-4OHT induction is challenging with standard stereo microscopy due to the slow reporter accumulation in *ubi:Switch* (**Fig. 3C,D**). We did not observe any switched pectoral fin cells, further supporting our conclusion that *hsp70l*:*Switch* provides more thorough recombination reporting than *ubi:Switch*. Together, these results underscore that the original *ubi:creERT2* transgenic (*cz1702Tg, ZDB-ALT-110121-1*; Mosimann et al., 2011) features diminished recombination potency at later developmental stages.

### Modifying *EGFP* in *ubi:Switch* does not alter recombination efficiency

CRISPR-Cas9-mediated genome editing has become a standard tool to generate zebrafish mutants. Cas9 can also be harnessed to target transgenic insertions to mutate regulatory elements and transgenic cargo such as fluorescent protein ORFs. Modifying efficient *lox*-based *Switch* transgenes provides an accessible method to alter the properties of reporter transgenes, yet the impact of such modifications to *lox* site recombination and expression efficiency remains unexplored. *ubi:Switch* expresses EGFP by default, resulting in ubiquitous green fluorescence that prohibits combining the reporter with GFP-based transgenic reporters (**Fig. 1A**). To evaluate the impact of *in situ* modification of a *Switch* locus in zebrafish, we isolated a CRISPR-induced 4 base pair frameshift mutation (*Δ4*) in the *EGFP* ORF, resulting in a premature stop codon (**Fig. 4**). We derived the *ubi:Switch*-based transgenic strain *Tg(ubi:loxP-delta-EGFP-loxP_mCherry)* (*ubi:delta-EGFP* for short) and compared this new transgene to the original *ubi:Switch* (**Fig. 4**).

**Figure 4.**
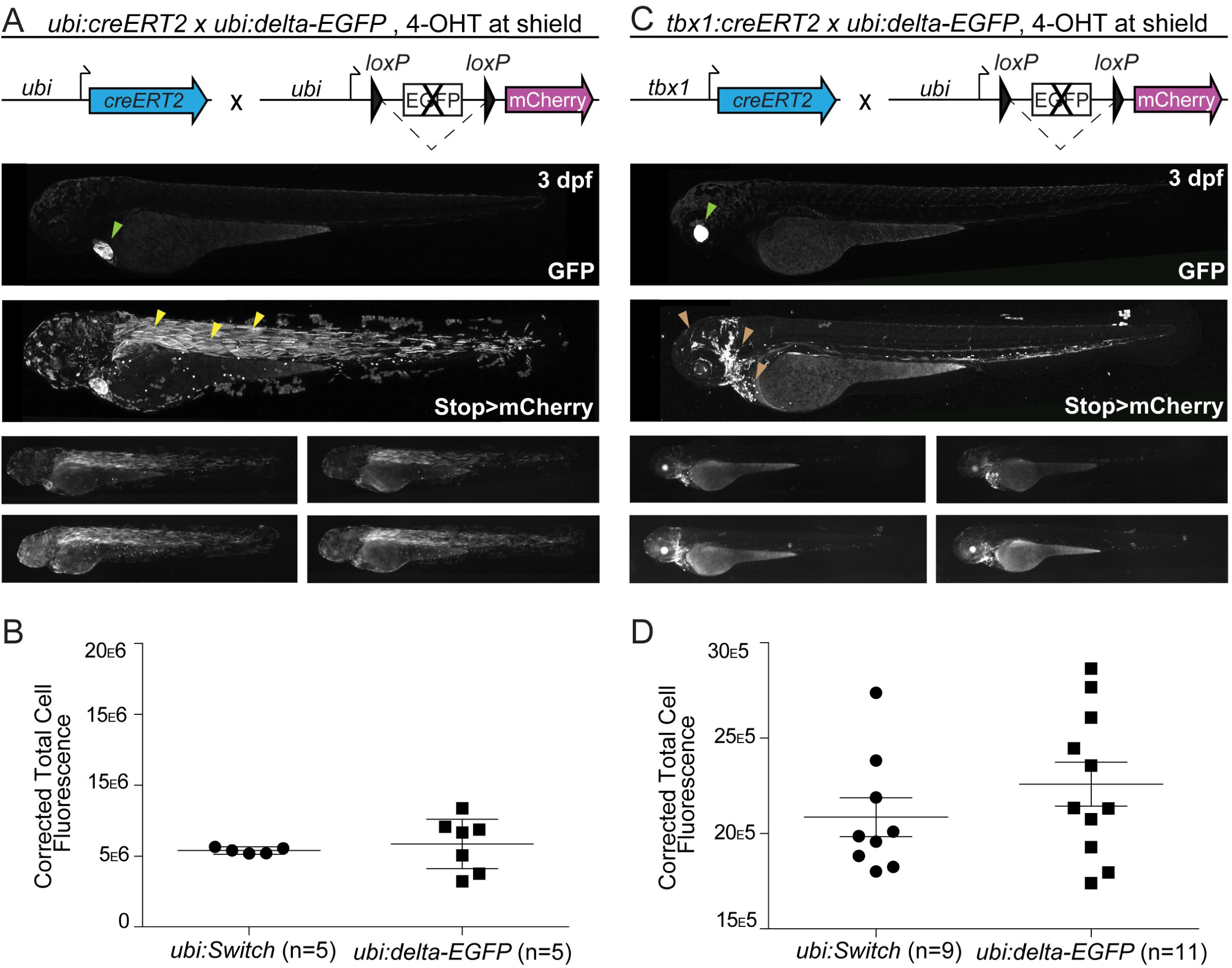
Modifying *EGFP* in *ubi:Switch in situ* does not alter recombination efficiency. (**A**,**B**) *ubi:creERT2* and (**C**,**D**) *tbx1:creERT2* crossed to *ubi:delta-EGFP* that bases on *ubi:Switch* with a disrupted EGFP cassette, induced with 4-OHT at shield stage, and imaged laterally at 3dpf plus corrected total cell fluorescence for quantification (**B**,**D**). One representative confocal image, and four representative stereoscopic images are presented here (**A**,**C**). *ubi:creERT2* shows high recombination efficiency with preferential switching in somitic muscle (yellow arrowheads) (**A**). No significant difference is observed when *ubi:creERT2* is combined with either *ubi:Switch* (STDEV.S=2.06E+05) or *ubi:delta-EGFP* (STDEV.S=1.89E+06) (Mann-Whitney, p>0.05) (**B**). *tbx1:creERT2* switching in the ventricular cardiomyocytes, pharyngeal arches, cranial vasculature, head muscles and cartilage, and hatching gland with *ubi:delta-EGFP* (brown arrowheads) (**C**). No significant difference in recombination efficiency is observed between *tbx1:creERT2* crossed to *ubi:Switch* (STDEV.S=3.05E+05) or *ubi:delta-EGFP* (STDEV.S=3.80E+05) (Mann-Whitney, p>0.05) (**D**). Note visibility of screening markers (*myl7:EGFP for ubi:creERT2; cryaa:Venus for tbx1:creERT2*) when EGFP ORF is removed in *ubi:delta-EGFP* (lime arrowhead) (**A**,**C**).

When crossed to *ubi:creERT2* and treated with 4-OHT at shield stage, *ubi:delta-EGFP* showed preferential switching in the zebrafish skeletal muscle at 3 dpf (**Fig. 4A**). To quantify the switching efficiencies of *ubi:Switch* and *ubi:delta-EGFP*, we performed whole-mount lateral view measurements of fluorescence intensity. These measurements did not reveal any significant differences in recombination pattern, mosaicism, or mCherry intensity at these experimental time points (n = 5, **Fig. 4B**). When crossed to *tbx1:creERT2, ubi:delta-EGFP* showed labeling in the ventricular cardiomyocytes, pharyngeal arches, cranial vasculature, head muscles and cartilage, and hatching gland at 3 dpf (**Fig. 4C**), with no significant difference to *tbx1:creERT2, ubi:Switch* (n = 9-11, **Fig. 4D**). Although neither tissue-specific creERT2 driver line showed significant differences in fluorescence measurements between *ubi:Switch* and *ubi:delta-EGFP, ubi:delta-EGFP* showed higher variability, as indicated by standard deviation (**Fig. 4B,D**). From this limited experimental paradigm, we conclude that *ubi:delta-EGFP* broadly preserves the recombination efficiency of *ubi:Switch* after modification of the fluorescent ORF.

### Transgene mapping reveals genomic features at *Switch* reporter integrations

The rational selection of suitable locations for universal, inert, and permissive genomic integration spots for transgenes is highly desirable to minimize position effects. While the majority of zebrafish transgenesis has been performed with random integration methods including Tol2 and ISce-I, CRISPR-Cas9 now provides a first means for targeted transgene integrations. However, neither rational design principles for safe harbor sites or long term-validated loci for functional transgene integration, especially of functional *lox*-based *Switch* reporters, have been reported in zebrafish (Carney and Mosimann, 2018). Our data above established that the *Tol2*-based transgenes *hsp70l:Switch* and *ubi:Switch* integrated into loci that are highly permissive to *loxP* cassette recombination and can be maintained for over a dozen generations (Felker et al., 2018; Mosimann et al., 2011), possibly providing suitable genomic coordinates for universal transgene landing sites in the genome.

We used Thermal Asymmetric Interlaced (TAIL)-PCR to map the genomic integration loci of the *ubi:Switch* and *hsp70l:Switch* transgenes (Liu and Whittier, 1995). We sequenced the resulting PCR products, charted their genomic position by BLAST, and confirmed the transgene integrations with independent primers. The results of our transgene mapping are depicted in **Fig. 5A,B**. The *ubi:Switch* transgene integrated at *chr14:32826253*, proximal and upstream of the gene *inppl1b* and distal and upstream of the gene *arr3b* (**Fig. 5A**); despite this proximity to gene bodies, homozygous *ubi:Switch* zebrafish are viable with no overt morphological phenotypes. The *hsp70l:Switch* transgene integrated at *chr24:26386749*, placing it 122 bp from the transcription start of an annotated transcript of the gene *skilb*, which itself is annotated to be within the gene body of the gene *si:ch211-230g15*.*5* (**Fig. 5B**). While no loss-of-function alleles for *skilb* or *si:ch211-230g15*.*5* have been reported to our knowledge, we found also *hsp70l:Switch* to be homozygous viable and fertile with no discernible phenotype.

**Figure 5.**
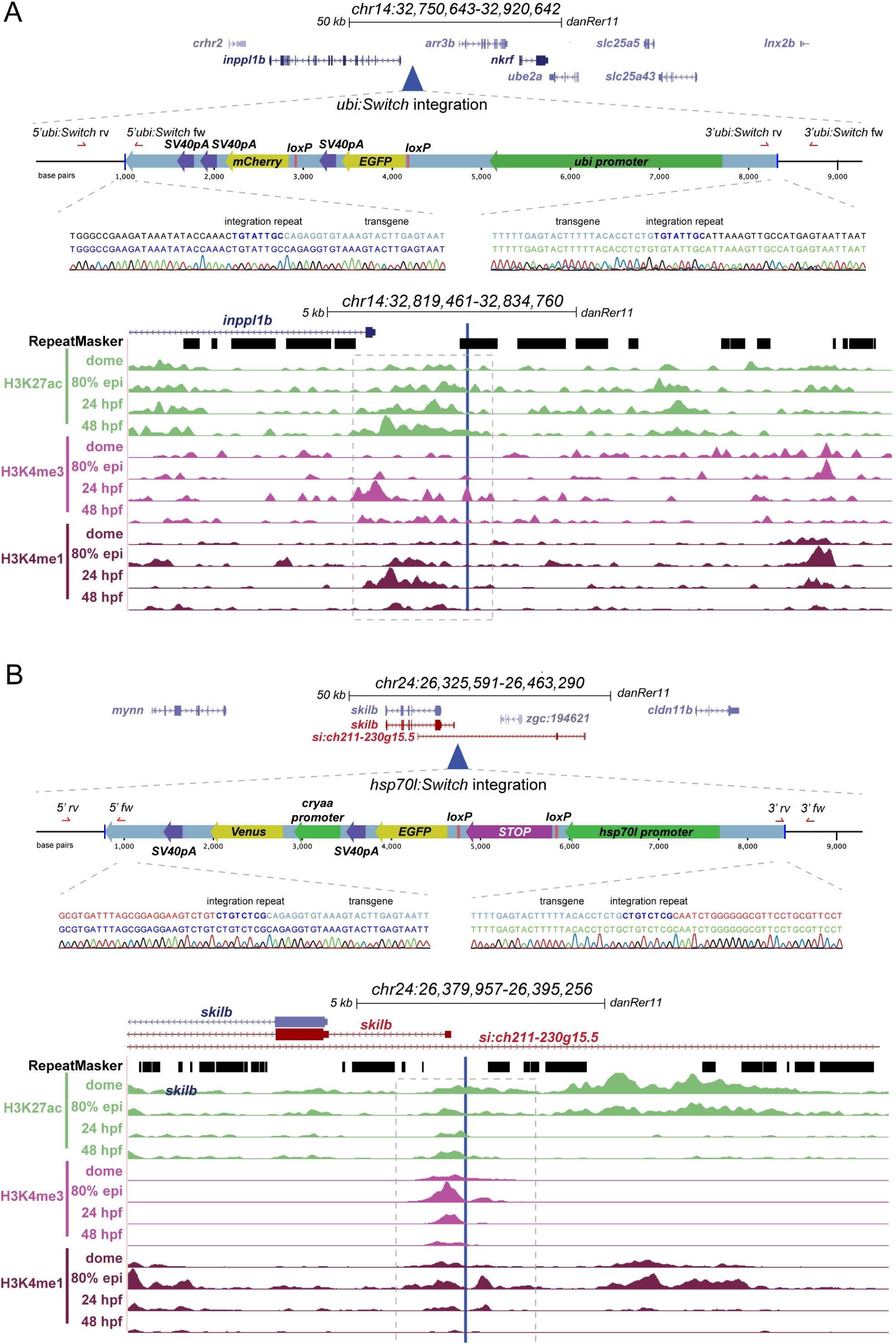
Transgene mapping reveals genomic features at the *ubi:Switch* and *hsp70l:Switch* loci. (**A**,**B**) Genomic integration sites of *ubi:Switch* and *hsp70l:Switch* determined by TAIL-PCR. For each transgene, the integration site is displayed in UCSC genome browser, a schematic of the transgene is depicted, the integration site is aligned to RepeatMasker track, and ChIP-seq tracks for active chromatin marks at key developmental stages. Sequencing reads for 5’ and 3’ transgene/genome boundaries are provided for *ubi:Switch* and *hsp70l:Switch* integration sites. *ubi:Switch* and *hsp70l:Switch* integration site loci contain peaks for all three chromatin modification signatures (H3K27ac, H3K4me3, H3K4me1, dashed boxes) (**A**,**B**).

In contrast to the *ubi:Switch* and *hsp70l:Switch* transgenic lines, we observed significantly less recombination with both the *actb2:Stop-DsRed* and the *actb2:BFP-DsRed* transgenic lines under the same experimental conditions (**Fig. 1A-D**). We again used TAIL-PCR to map the genomic integration loci of the *actb2:Stop-DsRed* and *actb2:BFP-DsRed* transgenes (**Fig. 6A,B**). The *actb2:Stop-DsRed* transgene integrated at *chr7:5941511*, within an intron of two distinct annotated genes: *dusp19b* and *si:dkey-23a13*.*11* (**Fig. 6A**). The *actb2:BFP-DsRed* transgene integrated at *chr19:25353744*, within the intron of the gene *glcci1a* (**Fig. 6B**).

**Figure 6.**
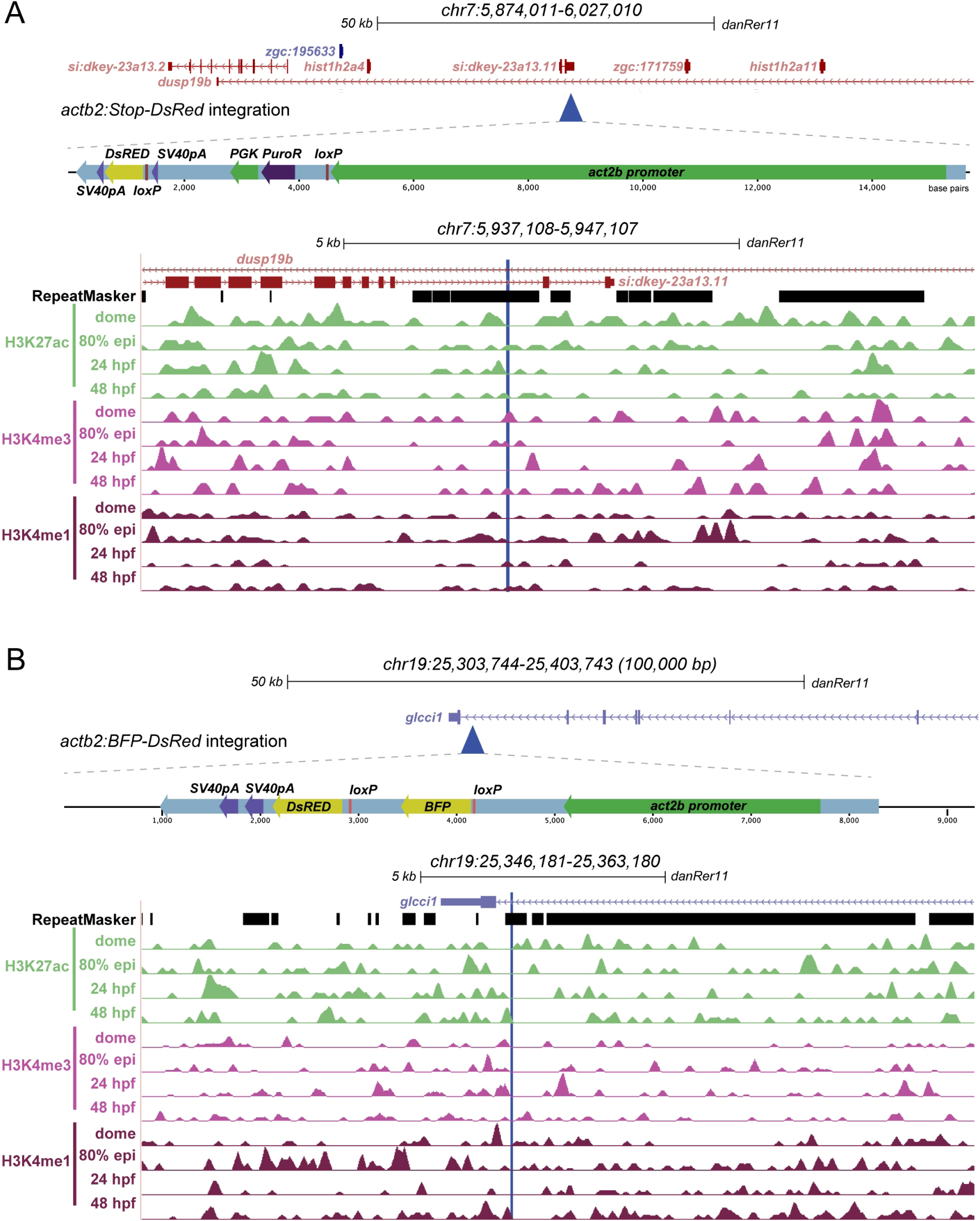
Transgene mapping reveals genomic features of the tested *actb2*-based *Switch* transgenes. (**A**,**B**) Genomic integration sites of *actb2-Stop-DsRed* and *actb2:BFP-DsRed* determined by TAIL-PCR. For each transgene, the integration site is displayed in UCSC genome browser, a schematic of the transgene is depicted, the integration site is aligned to RepeatMasker track, and ChIP-seq tracks for active chromatin marks at key developmental stages (**A**,**B**). Note the intronic integration sites for both transgenes and the absence of concise active chromatin marks compared to *ubi:Switch* and *hsp70l:Switch*.

To gain insight into what genomic features might influence *Switch* transgene recombination efficiency, we compared the integration sites of the mapped transgenes. All four transgenes mapped within or near protein-coding loci, with *ubi:Switch, actb2:Stop-DsRed, and actb2:BFP-DsRed* landing in highly repetitive regions (**Fig. 5A, 6A, 6B**). While *ubi:Switch* and *hsp70l:Switch* mapped to intergenic regions (*hsp70l:Switch* may lie within one possible gene body annotation) (**Fig. 5A,B**), both *actb2 Switch* transgenes mapped within introns of annotated and studied genes (*dusp19b* and *glcc1a*) (**Fig. 6A,B**). Using publicly available ChIP-seq datasets (Bogdanovic et al., 2012), we explored whether distinct histone marks at the integration loci of these individual *Switch* transgenic reporters may associate with high recombination. Distinct from surrounding regions, the *ubi:Switch* integration site showed a series of peaks for H3K27ac, H3K4me3, and H3K4me1 that represent open, active chromatin between 80% epiboly and 48 hpf (**Fig. 5A**). The immediate vicinity of the *hsp70l:Switch* integration locus featured signal for all histone marks between dome stage and 48 hpf (**Fig. 5B**). In contrast, the integration sites of the *actb2*-based *Switch* lines did not show active histone marks at any developmental stage we plotted (**Fig. 6A,B**). Based on a small sample size of four transgene insertions of different recombination qualities, these observations suggest that higher recombination efficiency might correspond with transgene integration in native areas containing active chromatin that may be primed for transcription. In addition, our mapping provides the genomic coordinates of the *ubi:Switch* and *hsp70l:Switch* reporters for potential future modifications and knock-ins of alternative reporters or *lox*-based *Switch* transgenes at these validated loci.

## Discussion

Transgenic experiments in zebrafish depend on reliable, reproducible reagents including single-copy insertion transgenes with predictable expression patterns in space and time. Cre/*lox* experiments in particular depend on efficient *lox*-based *Switch* reporters, the generation of which has so far been serendipitous due to favorable Tol2-based random transgene integrations. Here, we have documented the Cre-dependent recombination or switching efficiencies of four established *lox-*based *Switch* reporters with both ubiquitous and tissue-specific CreERT2 driver lines at two different developmental time points. Further, our results suggest two long-term validated genomic loci as potential safe harbor sites suitable for *lox*-based *Switch* reporters based on their reproducible performance over generations and across labs.

While seemingly straight-forward given access to suitable transgenic lines, Cre/*lox-*based lineage tracing studies require careful consideration to ensure successful experimental outcomes (Carney and Mosimann, 2018). The specific combination of fluorophores in the *lox*-based reporter may impede visualization if used in parallel with other desired transgenic reporter lines, such as when seeking to define switched cells with a tissue-specific reporter in parallel. Our conversion of *ubi:Switch* to *ubi:delta-EGFP* shows that modulating functional *Switch* transgenics *in situ* by small edits is feasible without drastically perturbing their properties (**Fig. 4**). Disruption of the *EGFP* ORF in *ubi:delta-Switch* facilitates the visualization of screening markers (*myl7:EGFP, cryaa:Venus*, etc.) and enables the use of this line in combination with commonly used EGFP/GFP-based reporter lines.

In Cre/*lox*-based lineage labeling, the onset of fluorescent reporter expression following *loxP* recombination dictates the earliest possible analysis timepoints. As documented in previous work, *ubi*-driven transgenes feature a latency in reporter expression, restricting detectable visualization and imaging to 24 hours or more after Cre-based recombination (**Fig. 3C,D**) (Carney and Mosimann, 2018; Chen et al., 2017; Felker et al., 2016; Mosimann et al., 2011). We note that, even with extended exposure length for imaging, visualizing mCherry fluorescence 24 hours post-4-OHT induction, remains challenging using standard stereo microscopy (**Fig. 3C,D**). Consequently, *ubi:Switch* may not be the preferred choice of *lox* reporter for short-term trace experiments (i.e. analysis within hours following first Cre activity), unless *in situ* hybridization for *mCherry* mRNA is used as a readout, rather than mCherry fluorescence. However, we here document that a simple heat shock prior to imaging can boost the reporter activity of *ubi:Switch* (**Fig. 1B**); if this pragmatic way to facilitate reporter detection with this transgene is based in heat shock elements in the zebrafish *ubi* promoter or based on other effects warrants further analyses. In contrast, the *hsp70l:Switch* transgenic line bypasses long latency in fluorescent reporter expression, yielding bright EGFP expression within 2 hours post heat-shock (**Fig. 1B, Fig. 2A,B**). As a potential caveat, heat-shock efficiency has not been extensively tested in adult *hsp70l:Switch* zebrafish; nonetheless the heat-shock system has been successfully used in adults with *Tg(hsp70:loxP-DsRed*-*Stop-loxP-EGFP)*^*tud107*^, as well as other *hsp70l*-based transgenic lines (Duszynski et al., 2011; Duszynski et al., 2013; Felker et al., 2018; Kroehne et al., 2011; Labonty et al., 2017; Pinzon-Olejua et al., 2017; Trompouki et al., 2011).

*ubi:creERT2* provides a versatile tool to test new *lox*-based *Switch* lines and to establish 4-OHT regimens (Carney and Mosimann, 2018; Felker et al., 2016; Mosimann et al., 2011). In our assays, inducing CreERT2 activity with 4-OHT during gastrulation leads to ubiquitous recombination in all tested *Switch* reporters (**Fig. 1**). When combined with *ubi:Switch*, we noted increased mCherry signal in the heart relative to other tissues, possibly a consequence of the heart-specific *myl7:EGFP* transgenesis marker of *ubi:creERT2* that could influence creERT2 expression (Huang et al., 2003; Kwan et al., 2007). Nonetheless, CreERT2 activity from *ubi:creERT2* is drastically reduced by 48 hpf despite substantial evidence that the *ubi* regulatory element remains active throughout all stages of zebrafish development (Mosimann et al., 2011). Our data here adds to the notion that this diminished activity is due to the particular *ubi:creERT2* transgene insertion and not due to issues with 4-OHT uptake of the zebrafish embryo and larva: we have documented efficient switching with *drl:creERT2* when induced at 48 hpf, confirming 10 μM 4-OHT penetration in superficial tissues such as fin fibroblasts and blood (**Fig. 3C,D**). Efficient *loxP* recombination has also been documented through labeling of oligodendrocytes at 6 dpf following a 5 μM 4-OHT induction at 5 dpf *Tg(mbpa:mCherry-T2A-CreERT2)*, and in hepatocytes in 1-2 year old zebrafish livers following 2 μM 4-OHT induction from 5-7 dpf and 10-11 dpf (*tp1:creERT2)* (Pinzon-Olejua et al., 2017; Zhang et al., 2021). Labeled liver hepatocytes were observed at 10 weeks of age using *TRE:creERT2* following 1 μM 4-OHT induction at 5 weeks for 3 consecutive days, however no description of 4-OHT delivery was described (Li et al., 2019). The maintained integration of *ubi:creERT2* is therefore best applied for basic tests and at early developmental stages to recombine *lox*-based *Switch* reporters.

Our observations here present the case that, ideally, multiple independent *Switch* lines should be used with individual Cre drivers to confirm findings and avoid biasing lineage information as not all *Switch* lines are created equal. *lox*-based *Switch* reporters are highly sensitive to position effects, possibly arising from differential chromatin accessibility across independent transgenic insertions (Carney and Mosimann, 2018). Our transgene mapping provides first details of the chromatin environment and dynamics at the integration sites of the four transgenes tested here. From this limited sample size, the well-recombining *hsp70l:Switch* stands out as having integrated adjacent to a region with native H3K4me1, K3K4me3, and H3K27Ac marks indicative of open, active chromatin from early developmental stages (Bogdanovic et al., 2012) (**Fig. 5**). The *ubi:Switch* integration locus showed H3K27Ac marks across developmental timepoints, yet lacks H3K4 methylation marks. In contrast, the two less Cre-responsive *actb2*-based *Switch* lines that integrated into introns show no consistent histone marks associated with open chromatin (**Fig. 6**). Nonetheless, all observed correlation with open histone marks is based on native genome context without inserted transgene, and how the Tol2-based integrations themselves affect the chromatin context remains unknown. Our mapping further uncovered that while *hsp70l:Switch* did not integrate within a repetitive region, *ubi:Switch, actb2:BFP-DsRed*, and *actb2:Stop-DsRed* integrated within genomic repeat sequences of different classes (**Fig. 5**,**6**). These first insights should encourage future mapping of Tol2 transgene insertions towards identifying features that support consistent transgene expression also beyond *Switch* lines.

So-called safe harbor sites to integrate transgenes provide a key tool for reproducible transgene deployment in any given model. Previously generated transgene integrations for repeated insertion based on the phiC31 recombinase system have either shown weak to medium expression levels, have not been validated for *lox*-based recombination, or have not been maintained beyond proof-of-principle (Bhatia et al., 2021; Carney and Mosimann, 2018; Hu et al., 2011; Lister, 2010; Lu et al., 2011; Mosimann et al., 2013; Roberts et al., 2014). Consequently, versatile and validated safe harbor sites are currently missing in zebrafish, rendering Tol2-or ISce-I-based random integration and subsequent screening for functional single-copy transgenes labor-intensive. Recent work has reported the Tol2-based generation of a phiC31-targetable *attB* integration site called *SHH-SBE2* that is suitable for gene-regulatory element analysis through reproducible transgenesis into the same locus (Bhatia et al., 2021); while promising, if *SHH-SBE2* shows long-term stability and is amenable to *lox*-based *Switch* reporters warrants further investigation. Retained switching efficiency over generations is a highly desirable feature of *lox*-based *Switch* reporter transgenics. CRISPR-Cas9-based knockin would provide means to target transgene insertion into suitable loci (Auer et al., 2014; Kesavan et al., 2018; Kimura et al., 2014; Prykhozhij et al., 2018; Shin et al., 2014); nonetheless, little predictive information is available as to what genomic loci provide safe harbor sites.

In mammalian systems, the serendipitously discovered *Hipp11* (*H11*) locus is widely used for reproducible transgenesis (Hippenmeyer et al., 2010; Tasic et al., 2011; Zhu et al., 2014). Our data argues that, in zebrafish, functional and stable Tol2 insertions of *lox*-based *Switch* transgenics indicate potential safe harbor sites that are also suitable for recombinase-sensitive cassettes. *ubi:Switch* and *hsp70l:Switch* have been maintained as functional lines in numerous laboratories since their isolation in 2009 (Mosimann et al., 2011) and 2016 (Felker et al., 2018), respectively. Considering the high recombination efficiency, and retained activity over generations of both *ubi:Switch* and *hsp70l:Switch* transgenes, we propose these loci as validated sites for future transgenic work. A possible approach involves CRISPR-Cas9-based removal of either *Switch* transgene and integration of a phiC31-targeted landing site, such as by targeting the Tol2 transposon arms 5’ and 3’ of the respective transgene and screening for lost reporter expression. Altogether, our observations argue that mapping of well-working Tol2 transgene insertions, especially of *lox*-based *Switch* reporters, should become more widespread in the zebrafish community to inform about suitable transgene insertion sites with validated activity.

## Acknowledgements

We thank Christine Archer and Molly Waters for zebrafish husbandry support, Dr. Oscar Ruiz for support with TAIL-PCR reagents, Dr. Caleb Doll for input on imaging, and all members of the Mosimann lab for input on the manuscript.

## Author Contributions

R.L.L., C.L.K., F.W.R performed the lineage-tracing imaging and analysis. R.L.L., and S.N. performed the transgene mapping. J.K.-R. assembled the chromatin modification signature ChIP-seq tracks in UCSC genome browser. R.L.L. and C.M. wrote the manuscript with contributions from all authors. C.L.K. created the figures. A.J.A made the *ubi:delta-EGFP* switch line under D.M.P.’s supervision. A.B. and C.M. conceptualized the study and supervised.

## Materials and Methods

### Zebrafish Husbandry and Procedures

Animal care and procedures were carried out in accordance with the veterinary office of the IACUC of the University of Colorado School of Medicine (protocol #00979), Aurora, CO, USA. Females from the included *Switch* lines were crossed with male *[regulatory-element]:creERT2* transgenic zebrafish using dividers and embryos were incubated at 28.5° C in E3 medium.

### Drug Administration and Heat shock Protocol

Activity of CreERT2 was induced with 10 μM final concentration of (Z)-4-Hydroxytamoxifen (Sigma Aldrich, St. Louis, MO, USA, H7904, abbreviated as 4-OHT) in E3. 4-OHT stock is stored at −20°C in the dark as 10mM single-use aliquots dissolved in DMSO and used within 2 months of dissolving. Prior to administration, the 4-OHT aliquots were incubated at 65°C for 10 minutes and vortexed. For shield stage treatment, 4-OHT was administered overnight and then replaced with N-Phenylthiourea (Sigma Aldrich, P7629, abbreviated as PTU) at a final concentration of 200 μM in DMSO embryo medium each morning to inhibit melanogenesis. For embryos treated with 4-OHT at 2 dpf, PTU was administered at 24 hpf, 4-OHT/PTU was administered at 48hpf, replaced with PTU at 56 hpf and refreshed the next morning prior to imaging. *Tg(hsp70l:Switch)* embryos were heat-shocked for 1 hour at 37°C in a water bath. Prior to heat shock embryos were dechorionated and transferred to a glass vial with E3 medium. Embryos were imaged 2 hours (stereo microscope) and 3 hours (confocal microscope) later.

The *ubi:delta-EGFP* transgenic line was made using the following sgRNA to target the *EGFP* ORF in *ubi:Switch*: *5’-GAGCTGGACGGCGACGTAAA-3’*. The mutant allele harbors a 4 bp deletion resulting in a frameshift mutation beginning at *Y40A*, and premature stop at *Stop42K. ubi:delta-EGFP* was generated by injecting *in vitro*-transcribed sg RNA and recombinant Cas9 protein (PNA Bio) into *ubi:Switch* zygotes (Burger et al., 2016; Hwang et al., 2013; Shah et al., 2015). Resulting F0s were screened for mosaic EGFP expression and a single, highly mosaic individual was selected to establish the line. Mutagenesis was confirmed by Sanger sequencing the PCR product amplified with Fw *5’-TTTAACATGGGAGAAGTGCAAAA-3’*; Rev *5’-GTCGTCCTTGAAGAAGATGGTG-3’*.

### Imaging

Embryos were anesthetized at 3dpf with 0.016% Tricaine-S (MS-222, Pentair Aquatic Ecosystems, Apopka, FL, USA, NC0342409) in E3 embryo medium. Basic fluorescence imaging was performed on a Leica M205FA with a DFC450 C camera. Laser scanning confocal microscopy was performed on a Zeiss LSM880 following embedding in E3 with 1% low-melting-point agarose (Sigma Aldrich, A9045) on glass bottom culture dishes (Greiner Bio-One, Kremsmunster, Austria, 627861). Images were collected with a x10/0.8 air-objective lens with all channels captured sequentially with maximum speed in bidirectional mode, with the range of detection adjusted to avoid overlap between channels. Maximum projections of acquired Z-stacks were made using ImageJ/Fiji (Schindelin et al., 2012) and cropped and rotated using Adobe Photoshop 2021.

### Quantifications

For CTCF (corrected total cell fluorescence) calculations, lateral images were taking using the stereo microscope and then processed using ImageJ (Felker et al., 2016; Schindelin et al., 2012). Whole embryos (*ubi:creERT2*) or anterior half (*tbx1:creERT2*) were traced using the magic wand tool and then measured. For every picture, the CTCF (corrected total cell fluorescence) was calculated by the formula ‘Integrated density whole–(area whole embryo x mean fluorescence background)’. This formula is loosely based on a method described for calculating cell-fluorescence (Burgess et al., 2010).

For ISV quantification lateral view confocal z-stacks were taken and then max projections were generated using imageJ. %ISV (Intersomitic vessels) were manually counted and calculated by dividing total switched ISV/total ISV (Felker et al., 2016).

For head vessel and muscle quantification ventral view confocal z-stacks were taken and then max projections were generated using imageJ. Head vessels/muscles were manually counted using *fli1a:EGFP* embryos stained with the MF20 antibody as a vessel/muscle control reference.

### Statistics

Unpaired non-parametric (Mann-Whitney) two-tailed t-test was done to compare the scores between two groups. For analyses with more than two groups, 1-Way ANOVA] was performed to compare the scores between the groups. Adjusted p-values after multiple tests correction are reported and significance was set at p < 0.05.

### MF20 immunostaining

*fli1a:EGFP* zebrafish (Lawson and Weinstein, 2002) were dechorionated and fixed at 3 dpf with 4% formaldehyde, 0.1% Triton-X100 in PEM (0.1 M PIPES, 2 mM MgSO4, and 1 mM EDTA) overnight at 4°C. The next day, embryos were washed in 1x PBS with 0.1% Triton-X100 and 0.1% BSA, permeabilized for 1 hour at RT with ProtK diluted in PBS to a final concentration of 5ug/ul and next for 30 min in 1x PBS with 0.5% Triton-X100 for 30 min. After washing, embryos were blocked in PBS + 1% goat serum and 0.1% Triton-X100 for 2 hours at room temperature and incubated overnight at 4°C with primary antibody MF20 (DSHB, antibody ID# AB_2147781) 1:50 diluted in blocking buffer. The 3rd day, embryos were washed 3 times 10 minutes in 1x PBS with 0.1% Triton-X100 and 0.1% BSA and incubated overnight at 4°C with the secondary antibody goat-anti-mouse A568 1:500 (Abcam, antibody ID# ab175473). On the 4th day, embryos were washed 3x 30 min with 1x PBS with 0.1% Triton-X100 and 0.1% BSA and kept at 4° C in Vectashield with DAPI diluted 1:5 in the same wash buffer until imaging.

### TAIL-PCR

Protocol based on previous work (Liu and Whittier, 1995; Mosimann et al., 2013):

1. Extract DNA from fin clips or single embryos
2. Prepare primer mixtures: Combine 1.5 μM of each *Tol2*-specific primer with 10 μM of each AD primer.
3. Pipet primary PCR reaction: 1 μl Genomic DNA, 4 μl Primer mix (primary + random), 2 μl 10x Polymerase Buffer, 2 μl dNTPs (2 mM), 0.2 μl Polymerase (Expand™ High Fidelity PCR System, Roche), 11.3 μl ddH2O.
4. Perform *primary* PCR step (See PCR conditions below).
5. Dilute 2 μl of the primary PCR product in 25 μl ddH2O and add to the secondary reaction: 2 μl Diluted PCR product, 4 μl Primer mix (secondary + random), 2 μl 10x Polymerase Buffer, 2 μl dNTPs (2 mM), 0.2 μl Polymerase, 9.8 μl ddH2O.
6. Perform *secondary* PCR reaction (See PCR conditions below).
7. Dilute 2 μl of the secondary PCR product in 25 μl ddH2O and add to tertiary reaction. Pipette reaction analogous to the secondary reaction with tertiary primer and secondary product.
8. Perform *tertiary* PCR reaction (See PCR conditions below).
9. Compare secondary and tertiary PCR product on a 1.5% (w/v) agarose gel. If the amplification of the genomic region was successful, a slight band shift should occur between the secondary and the shorter tertiary product.
10. Excise bands and purify with QIAquick gel extraction kit (#NA1111-1KT).
11. Sequence purified PCR products.
12. Use BLAST (http://blast.ncbi.nlm.nih.gov/) to align to the zebrafish genome and identify 5’ and 3’ flanking regions.

The following gene-specific primers and random primers (AD3, 5, 6, 11) were used for TAIL-PCR:

*Tol2 5’1 forward GGGAAAATAGAATGAAGTGATCTCC*

Tol2 TIR 5’

*Tol2 5’2 forward GACTGTAAATAAAATTGTAAGGAG*

Tol2 TIR 5’

*Tol2 5’3 forward*

*CCCCAAAAATAATACTTAAGTACAG*

Tol2 TIR 5’

*Tol2 3’1 reverse CTCAAGTACAATTTTAATGGAGTAC*

Tol2 TIR 3’

*Tol2 3’2 reverse ACTCAAGTAAGATTCTAGCCAGA*

Tol2 TIR 3’

*Tol2 3’2 reverse CCTAAGTACTTGTACTTTCACTTG*

Tol2 TIR 3’

*AD-3 WGTGNAGNANCANAGA* Random

*AD-5 WCAGNTGWTNGTNCTG* Random

*AD-6 STTGNTASTNCTNTGC* Random

*AD-11 NCASGAWAGNCSWCAA* Random

Primers needed for the mapping of Tol2 integrations as described previously (Parinov et al., 2004): *W*: Weak base (*A* ort *T*), *S*: Strong base (*C* or *G*), *N*: Any base (*A, C, G, T*).

The following PCR conditions were used to TAIL-PCR:

Primary PCR reaction:

1. 2 min 94° C
2. 0.5 min 94° C
3. 1 min 62° C
4. 2.5 min 72° C
5. Repetition of steps 2-4 (5x)
6. 0.5 min 72° C
7. 3 min 25° C
8. Ramping 0.3°C/s to 72° C
9. 2.5 min 72° C
10. 10 s 94° C
11. 1 min 61° C
12. 2.5 min 72° C
13. 10 s 94° C

Secondary PCR reaction:

1. 10 s 94° C
2. 1 min 61° C
3. 2.5 min 72° C
4. 10 s 94° C
5. 1 min 61° C
6. 2.5 min 72° C
7. 10 s 94° C
8. 1 min 44° C
9. Ramping 1.5° C/s to 72° C
10. 2.5 min 72° C
11. Repetition of steps 1-10 (15x)
12. 5 min 72° C

Tertiary PCR reaction:

1. 0.25 min 94° C
2. 1 min 44° C
3. Ramping 1.5° C/s to 72° C
4. 2.5 min 72° C
5. Repetition of steps 1-4 (30x)
6. 5 min 72° C

### Transgene mapping confirmation

The following locus-specific primers were used to confirm the integration sites of *ubi:Switch* and *hsp70l:Switch*:

*5’ ubi:Switch Fw GGAGCATTCAGAGGTACC*

*5’ ubi:Switch Rev GACTGTAAATAAAATTGTAAGGAG*

*3’ ubi:Switch Fw CTCAAGTACAATTTTAATGGAGTAC*

*3’ ubi:Switch Rev GCTGTGAGACGATCAGGC*

*5’ hsp70l:Switch Fw GCATGACACGGCTAACCAAC*

*5’ hsp70l:Switch Rev GACTGTAAATAAAATTGTAAGGAG*

*3’ hsp70l:Switch Fw CCTAAGTACTTGTACTTTCACTTG*

*3’ hsp70l:Switch Rev TATCAGCACACACCTTTATCGC*

We were unable to fully confirm the integration sites for the *actb2* Switch transgenes with independent locus-specific primers. We confirmed the 5’ genomic border of the *actb2:BFP-DsRed* transgene, however the 3’ genomic border and both 5’ and 3’ genomic borders of *acbt2:Stop-DsRed* could not be confirmed.

---

**Figure S1:**
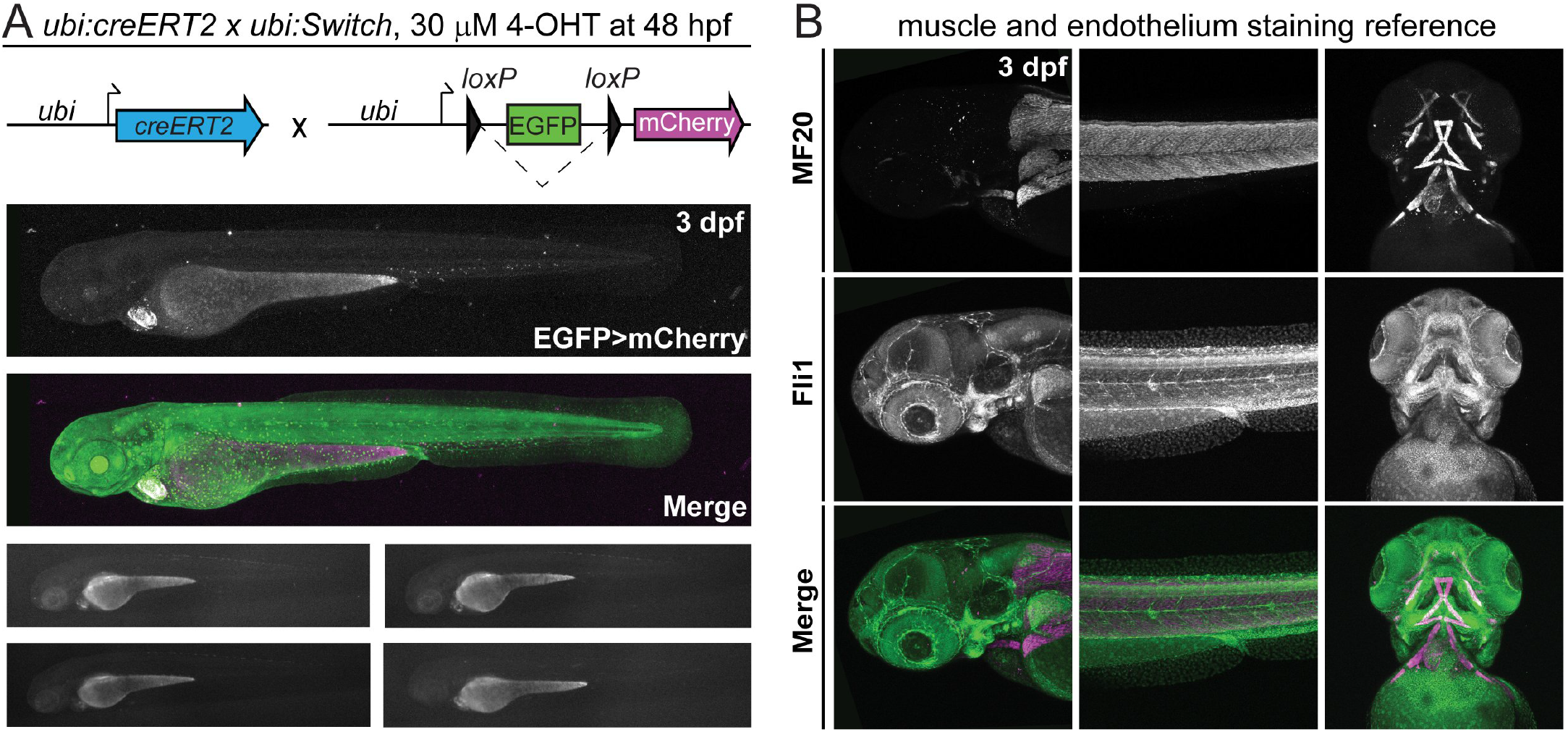
(**A**) *ubi:creERT2* crossed to *ubi:Switch* induced with 30μM 4-OHT at 48hpf, and imaged lateral at 3dpf. No increase in switching efficiency is observed with 30 μM as compared to 10 μM (**Fig. 3B**). (**B**) Max projections of z-stack confocal images of *fli1a:EGFP* transgenic zebrafish fixed at 3dpf and stained for muscles using an MF20 antibody to visualize vasculature and muscle anatomy in the zebrafish head at 3dpf.

